# A 2022 avian H5N1 influenza A virus from clade 2.3.4.4b attaches to and replicates better in human respiratory epithelium than a 2005 H5N1 virus from clade 2.3.2.1

**DOI:** 10.1101/2024.11.27.625596

**Authors:** Lisa Bauer, Lonneke Leijten, Matteo Iervolino, Varun Chopra, Laura van Dijk, Mark Power, Monique Spronken, Willemijn Rijnink, Mathis Funk, Rory D. de Vries, Mathilde Richard, Thijs Kuiken, Debby van Riel

## Abstract

**Background:** Highly pathogenic avian influenza (HPAI) H5 viruses of the A/Goose/Guangdong/1/1996 (GsGd) lineage pose significant global risks to wildlife, domestic animals, and humans. Recent cross-species transmission events to mammals, including humans, highlight this risk. Critical determinants for cross-species and intra-species transmission include the ability to attach to and replicate in respiratory epithelial cells. Although these factors have been studied for HPAI H5N1 viruses in the past, limited studies are available for currently circulating strains.

**Methods:** We compared level of adaptation to human respiratory tract of a HPAI H5N1 clade 2.3.4.4b (H5N1^2022^) virus with those of well characterized HPAI H5N1 clade 2.1.3.2 (H5N1^2005^) and seasonal H3N2^2003^ viruses by three methods. First, we compared pattern of virus attachment by virus histochemistry. Second, we compared efficiency of infection and replication, as well as innate immune responses in human respiratory epithelium *in vitro*. Lastly, we compared polymerase complex activity in a minigenome assay.

**Findings:** The H5N1^2022^ virus attached more abundantly to and replicated more efficiently in cells of the human respiratory tract compared to H5N1^2005^ and H3N2 viruses. This increased replication was not associated with an increased polymerase activity of H5N1^2022^ virus compared to H3N2^2003^ virus. The efficient replication of H5N1^2022^ virus infection induced a robust innate immune response almost comparable to H3N2^2003^.

**Interpretation:** The pattern of virus attachment and replication efficiency of a HPAI H5N1^2022^ virus resembled that of H3N2^2003^ virus more closely than a HPAI H5N1^2005^. This could contribute to an increased risk for both human infection and virus adaptations to humans.

**Funding:** The Netherlands Organization for Health Research and Development

**Research in context:** *Evidence before this study:* Highly pathogenic avian influenza (HPAI) H5 viruses of the A/Goose/Guangdong/1/1996 (GsGd) lineage (clade 2.3.4.4b) have the ability to spread to a wide range of domesticated and wild mammalian species, including humans. Cross species transmission and transmission among humans requires— among other factors—efficient infection of epithelial cells in the respiratory epithelium of the upper respiratory tract.

*Added value of this study:* In our study we show that a recent clade 2.3.4.4b HPAI H5N1 virus attached to and replicated more efficiently in respiratory epithelium than a clade 2.1.3.2 H5N1 virus that circulated in 2005.

*Implications of all the available data:* These data suggest that there might be an increased risk of human infections with the currently circulating 2.3.4.4b HPAI H5N1 viruses, which might facilitate opportunities for human adaptation.

## Introduction

In 1996, a highly pathogenic avian influenza (HPAI) H5N1 virus of the A/Goose/Guangdong/1/96 (GsGd) lineage emerged in domestic geese^1^. Over time, viruses of the GsGd lineage have spread worldwide in wild birds and poultry, causing significant economic losses in the poultry industry and occasionally spill-over into mammals including humans^2^. Since 2020, HPAI H5N1 clade 2.3.4.4b viruses of the GsGd lineage are circulating continuously in wild birds, leading to further intercontinental spread^3–6^, and causing mass mortality events in various wild birds^7,8^, a variety of free-living wild and marine mammals^9–12^, and farmed fur-bearing mammals such as foxes and mink^13,14^. Recently, a spill-over of HPAI H5N1 virus of clade 2.3.4.4b into dairy cows resulted in wide-spread circulation in the United States^15,16^. The extensive spread of H5N1 viruses in wild and domestic animals poses an increased potential threat for spill-over transmission to humans^17^, exemplified by the recent infection of dairy farm workers in the United States^18,19^.

Influenza A viruses crossing species barriers^20^ can result in sometimes devastating effects for instance by the human pandemic in 1918^21^, or the recent mass mortality event in sea lions in South America^11^. The ability of influenza A viruses to cross species barriers and spread within a new host species depends on viral and host factors^20^. For example, the receptor recognition of influenza A viruses, as well as the appropriate distribution of these receptors in the host are thought to be critical. Subsequent inter-species transmission is often associated with genetic changes in viral genes such as those coding for the surface glycoprotein haemagglutinin (HA) or encoding the subunits of the viral polymerase complex (PB1, PB2, PA)^22,23^.

The ability to attach to and replicate in tissues of the upper respiratory tract is critical for efficient infection and transmission of influenza A viruses, whereas attachment to and replication in cells in the lower respiratory tract is associated with its ability to cause severe respiratory disease^24–26^. Attachment of influenza A viruses to host cells is mediated by binding of the viral surface glycoprotein HA to sialic acid moieties of glycoproteins or glycolipids on the host cell surface. The dichotomy of receptor-binding preferences for avian and human influenza A viruses plays a crucial role in the host range of these viruses^27^. Simply stated, avian influenza A viruses prefer to bind to α2– 3-linked sialic acids present in the avian digestive tract^28^ as well as in the human lower respiratory tract^28,29^. In contrast, human influenza viruses preferentially bind to α2–6-linked sialic acids that are abundantly present in the human upper respiratory tract^24,29–31^. Consequently, for efficient infection of, and transmission among humans, influenza A viruses need to overcome the differences between avian and human receptor repertoires in order to attach to and replicate in cells in the human upper respiratory tract.

Currently circulating HPAI H5N1 clade 2.3.4.4b viruses have retained a preference for 2,3-linked sialic acids, based on glycan array studies^32–34^. In addition, transmission studies of H5N1 clade 2.3.4.4b viruses in ferrets, a model for virus interspecies transmission among humans, suggest inefficient airborne, but efficient contact transmission^34–37^. However, the ability of currently circulating HPAI H5N1 2.3.4.4b viruses to attach to and replicate in the human respiratory tract, and how this compares to older HPAI H5 virus clades and seasonal human influenza viruses, has not been studied.

Here, we compared the attachment pattern of a clade 2.3.4.4b H5N1 virus with a clade 2.1.3.2 H5N1 virus and a seasonal H3N2 virus in the human respiratory tract using virus histochemistry (VHC). Furthermore, we investigated the efficiency of infection and replication in primary nasal and tracheal/bronchial respiratory epithelium as well as the associated innate immune responses. Lastly, we determined if the observed differences in replication efficiency among the HPAI H5N1 viruses were associated with the polymerase complex activity.

## Methods

### Cell Culture

Madin-Darby canine kidney (MDCK) cells were cultured in Eagle’s Minimal Essential Medium (EMEM; Capricorn Scientific) supplemented with 10% fetal bovine serum (FBS; Sigma Aldrich), 100 IU/ml penicillin (Capricorn Scientific), 100 μg/ml streptomycin (Capricorn Scientific), 2 mM glutamine (Capricorn Scientific), 10 mM HEPES (Capricorn Scientific), and 0.1 mM nonessential amino acids (Capricorn Scientific). Human embryonic kidney (HEK293T) cells (ATCC) were cultured in Dulbecco’s Modified Essential Medium (DMEM; Capricorn Scientific) supplemented with 10% FBS (Sigma Aldrich), 2mM L-Glutamine (Capricorn Scientific) and 100U/ml penicillin/streptomycin (Capricorn Scientific). Cells were passaged at >90% confluence and routinely checked for the presence of mycoplasma.

### Human nasal respiratory epithelium

MucilAir tissues from a pool of donors were purchased from Epithelix Sàrl, Geneva, Switzerland. The upper airway epithelium was cultured at ALI in the medium provided by Epithelix according to the manufacturers protocol.

### Tracheal/bronchiolar Epithelium

Airway organoids at ALI were grown as described previously^16,38–41^. In short, bronchial AO were cultured in Matrigel (Corning) for ∼2 weeks, after which they were disrupted to single cells using TryplE (Gibco) and seeded on Rat Tail Collagen type I (Corning) coated transwells (Costar, Corning). After 2-4 days a monolayer had formed and transwells were put at ALI. AO cultures were differentiated for 4-6 weeks at ALI.

### Viruses

The HPAI H5N1 clade 2.3.4.4b virus isolate (A/Caspian gull/Netherlands/1/2022 [H5N1], referred to as H5N1^2022^ virus, EPI_ISL_16524246) was isolated from a Caspian gull (*Larus cachinnans*) and propagated three times in MDCK cells. The HPAI H5N1 virus from clade 2.1.3.2 (A/Indonesia/5/2005 ([H5N1], referred to as H5N1^2005^ virus) was isolated from a human patient^25,42^, and the virus was propagated once in embryonated chicken eggs and twice in MDCK cells. The human seasonal influenza A virus (A/Netherlands/213/2003 ([H3N2]; referred to as H3N2^2003^ virus) was isolated from a patient and the virus was propagated three times in MDCK cells.

### Generation of recombinant viruses for virus histochemistry

Recombinant viruses were generated as previously reported^43,44^. In short, recombinant viruses using 7 segments of the mouse-adapted influenza A virus strain A/Puerto Rico/8/1934 (PR8) and the HA segment of either H5 A/Indonesia/5/2005, H5 A/Caspian gull/Netherlands/1/2022 or H3 A/Netherlands/213/2003 was used. Site-directed mutagenesis was performed with the Pfu Ultra II Fusion HS DNA Polymerase (Agilent) and specific primers to remove the multibasic cleavage site (MBCS) which was replaced by the conserved H5 low pathogenic cleavage site. HEK 293T were transfected with the plasmids using the calcium phosphate–mediated transfection and a total of 40 μg of plasmid DNA was transfected. Approximately 16 hours after transfection, the cells were washed once with phosphate-buffered saline (PBS) and fresh media containing 2% FCS with 200–350 μg/mL *N*-tosyl-L-phenylalanine chloromethyl ketone (TPCK)-treated trypsin (Sigma-Aldrich) was added. Virus stocks were generated by inoculating MDCK cells s with dilutions of the supernatant harvested from the HEK293T cells 3 days post-transfection. The presence of the virus was confirmed by virus titration. Sequences from all plasmids and the HA genes of all virus stocks were confirmed with Sanger sequencing using the BigDye Terminator v3.1 Cycle Sequencing Kit (Applied Biosystems) and the 3500xL Genetic Analyzer (Applied Biosystems).

### Virus Titration

Virus titres were determined by endpoint dilution. Prior to titration, a confluent layer of MDCK cells was plated in 96-wells. 10-fold serial dilutions of supernatant in technical triplicate were prepared in infection medium (EMEM supplemented with 100 IU/ml penicillin, 100 μg/ml streptomycin, 2 mM glutamine, 1.5 mg/ml sodium bicarbonate, 10 mM HEPES, 1× (0.1 mM) nonessential amino acids, and 1 μg/μl tosylsulfonyl phenylalanyl chloromethyl ketone (TPCK)-treated trypsin (Sigma-Aldrich). Before adding the titrated supernatants, the MDCKs were washed once with plain EMEM medium supplemented with 100 IU/ml penicillin and 100 μg/ml streptomycin. Hundred µl of 10-fold serial dilution was used to inoculate MDCKs. After 1-hour incubation at 37°C, the supernatant was removed and 200µl fresh infection medium was added. Three days after infection, 25 µL of supernatant was mixed with 75 µL 0.25% turkey red blood cells (RBC) and incubated for 1 hour at 4°C and tested for agglutination. Infection titers were calculated according to the method of Spearman and Kärber and expressed as TCID50/ml^45^.

### Immunofluorescence Labelling

Cells on transwells were fixed by adding 10% formalin to the basolateral and apical compartment of the transwell for at least 30 minutes. Afterwards cells on the membrane were washed once with PBS and stored in PBS at 4°C. Next, cells on the transwell were permeabilized with PBS containing 1% Triton X-100 (Sigma) followed by a 30 minutes incubation in blocking solution consisting of PBS supplemented 0.5% Triton X-100 and 2% normal goat serum (Thermo Fisher Scientific) at room temperature. Primary antibodies diluted in blocking solution were used according to Table 1, and membranes incubated at room temperature for one hour. Membranes were washed three times with PBS and subsequently incubated with corresponding secondary antibodies, and Hoechst to visualize the nuclei, at room temperature for one hour. Stained membranes were washed three times in PBS and once in water, cut out of the transwell and mounted in ProLong Antifade Mountant (Invitrogen) on glass slides. Membranes were imaged using a Zeiss LSM 700 confocal microscope.

**Table 1.** Antibodies used for Immunofluorescence Microscopy.

| Antibodies | Cat nr | Final Concentration | Dilution | Manufacturer |
| --- | --- | --- | --- | --- |
| NP | EBS-I-047 | 2,0 µg/ml | 1:1000 | EVL laboratories |
| Acetylated-α-tubulin<br>Alexa Flour-647 | SC-23950 | 2,0 µg/ml | 1:100 | Santa Cruz Biotechnology |
| ZO-1<br>Alexa Flour-555 | 339100 | 5,0 µg/ml | 1:100 | Life Technologies/Invitrogen |
| Hoechst | 62249 | 2,0 µM | 1:10.000 | Life Technologies /Invitrogen |
| Donkey anti-Mouse Alexa Fluor-488 | A32766 | 2,0 µg/ml | 1:1000 | Life Technologies /Invitrogen |

### Multiplex bead assay for cytokine profiling

Cytokine concentrations in the supernatants were measured using the human antivirus response panel (13-plex) kit (LEGENDplex BioLegend). The kit was used according to the manufacturer’s manual with an additional fixing step. After adding the SA-PE and performing the washing steps, supernatant and the beads were fixed with formalin for 15 minutes at room temperature and washed twice with the provided wash buffer. This ensured that all pathogens were inactivated. The data were analysed with a Lyric flow cytometer (BD) and final analysis was performed using LEGENDplex analysis software v8.0.

### Virus preparation, inactivation and labelling for virus histochemistry

A confluent layer of MDCK cells were inoculated with influenza viruses. After 3 days supernatant was harvested and cleared by low-speed centrifugation. Cleared supernatants were subsequently centrifuged for 2 hours at 27,000 rpm in a SW28 rotor at 4 °C on a 0.5 ml layer of 60% sucrose. The lowermost 2.5 ml virus supernatant on top of the sucrose cushion was transferred to a 60-20 % sucrose gradient and centrifuged overnight at 39,000 rpm in a SW41 rotor at 4 °C. The virus fraction was harvested and diluted in PBS and centrifuged for 2 h at 27,000 rpm in a SW28 rotor at 4 °C, to deplete left-over sucrose. The virus pellet was resuspended in PBS and inactivated by dialysing against 0.1% formalin for 3 days at room temperature. Virus was labelled by mixing with an equal volume of 0.1 mg/ml of fluorescein isothiocyanate (FITC) (Sigma-Aldrich, Saint Louis, MO) in 0.5 M bicarbonate buffer (pH 9.5) for 1 hour at room temperature while constantly stirring. To lose all unbound FITC, labelled virus was dialysed against PBS, whereafter the HA titre was determined using the red blood cell hemagglutination assay as described in the virus titration section.

### Virus histochemistry on human respiratory tissues

Three-μm thick, formalin-fixed, paraffin-embedded human respiratory tract tissue sections were deparaffinized with xylene and rehydrated using graded ethanol. Approval from the Medical Research Ethics Committee was obtained to use human archival respiratory tract tissues for virus attachment studies (MEC-2011-129 and MEC-2009-128). Slides were incubated overnight at 4 °C with 100 µl FITC-labelled influenza virus (50-100 hemagglutination units). For visualization by light microscopy, FITC label was detected with a peroxidase labelled rabbit anti-FITC antibody (DAKOCytomation, Glostrup, Denmark). The signal was amplified with a tyramide signal amplification system (Perkin Elmer, Boston, MA) according to the manufacturer instructions. Peroxidase was revealed with 3-amino-9-ethyl-carbazole (AEC, Sigma-Aldrich) resulting in a bright red precipitate. Tissues were counterstained with hematoxylin and embedded in glycerol-gelatin (Merck, Whitehouse Station, NJ). Omission of the FITC-labelled virus was used as a negative control.

*Scoring of virus histochemistry:* Scoring was done as previously reported^24,46^ with slight adjustments. Tissue sections were scored using an Olympus BX51 microscope. The attachment of virus to the apical site of side of ciliated and non-ciliated respiratory epithelium, olfactory epithelium and pneumocytes was scored as follows: -, no attachment; +/-, attachment to rare or few cells (<10%); +, attachment to a moderate number of cells (10%-50%); ++, attachment to many cells (>50%).

### Immunohistochemistry on human airway epithelium

For detection of influenza A nucleoprotein in human nasal respiratory epithelium and tracheal/bronchiolar epithelium cultured on transwell membranes at ALI, cells were fixed with 10% buffered formalin for 30 minutes. The membranes were removed from the transwells and processed for paraffin embedding. Three-μm thick, paraffin-embedded sections were deparaffinized, rehydrated and antigen was retrieved in 0.1% protease for 10 minutes at 37°C. Endogenous peroxidase was blocked with 3% hydrogen peroxide. The slides were briefly washed with phosphate-buffered saline (PBS)/0.05% Tween 20 and incubated with mouse IgG2a-anti-Influenza A NP (Clone Hb65, American Type Culture Collection) 1/800 (2.5 µg/ml) in PBS/0.1% bovine serum albumin (BSA) or mouse IgG2a isotype control 1/200 (2.5 µg/ml) in PBS/0.1% BSA for 1 hour at room temperature. After washing, sections were incubated with horseradish peroxidase labelled goat-anti-mouse IgG2a (Serotec) 1/100 in PBS/0.1% BSA for 1 hour at room temperature. Peroxidase activity was revealed by incubating slides in AEC for 10 minutes, resulting in a bright red precipitate, followed by counterstaining with hematoxylin.

### Plasmids used for the minigenome assay

Plasmids in a PPI4 backbone containing the open reading frames of PB1, PB2, PA and NP to investigate the polymerase complex activity of H5N1^2005^ and H3N2^2003^ were kindly provided by Dirk Eggink. Plasmids in a pCAGGs backbone (kindly provided by Dr. A. Garcia-Sastre, Icahn School of Medicine, New York, USA) harbouring the open reading frames of PB1, PB2, PA and NP of H5N1^2008^ with either PB2 E627 or K627 polymorphism were previously described^47^

### Cloning of Caspian gull replication complex genes PB1, PB2, PA and NP in expression plasmids

The open reading frames of H5N1^2022^ PB1, PB2, PA and NP were cloned into the PPI4 backbone using In Fusion (Takara Bio) cloning. PCR fragments of the open reading frames were amplified using the primers listed in Table 2. For that viral RNA was isolated from virus stock using the Roche High Pure Viral Nucleic Acid kit according to the manufacturers protocol. SuperScript III kit was used to generate cDNA from RNA using random hexamer primers. cDNA was generated according to the manufacturers protocol protocol. cDNA was diluted 1:50 and 1 µl of diluted cDNA was used to amplify the viral genes. PCR product size was confirmed by running sample on 1% Agarose gel. DNA was extracted from the gel using Nucleospin Gel and PCR Clean-up (Machery-Nagel). The open reading frame of PA was not obtained, therefore the whole gene with overhangs for seamless cloning was ordered from Integrated DNA Technologies (IDT). The gBlock was reconstituted according to IDT instructions. The PPI4 backbone was digested with NotI and PstI and In-Fusion cloning was carried out using In-Fusion Snap Assembly kit (Takara Bio) according to the manufacturers protocol. Sequence of the plasmids were verified with the BigDye™ Terminator v3.1 Cycle Sequencing Kit (Applied Biosystems) and the 3500xL Genetic Analyzer (Applied Biosystems).

**Table 2.**
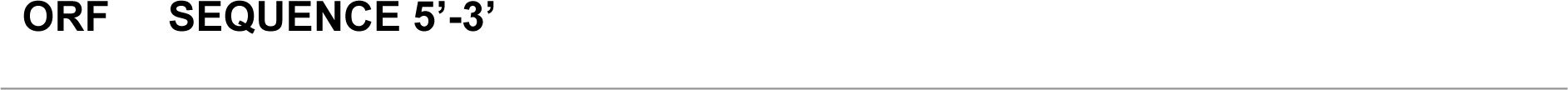
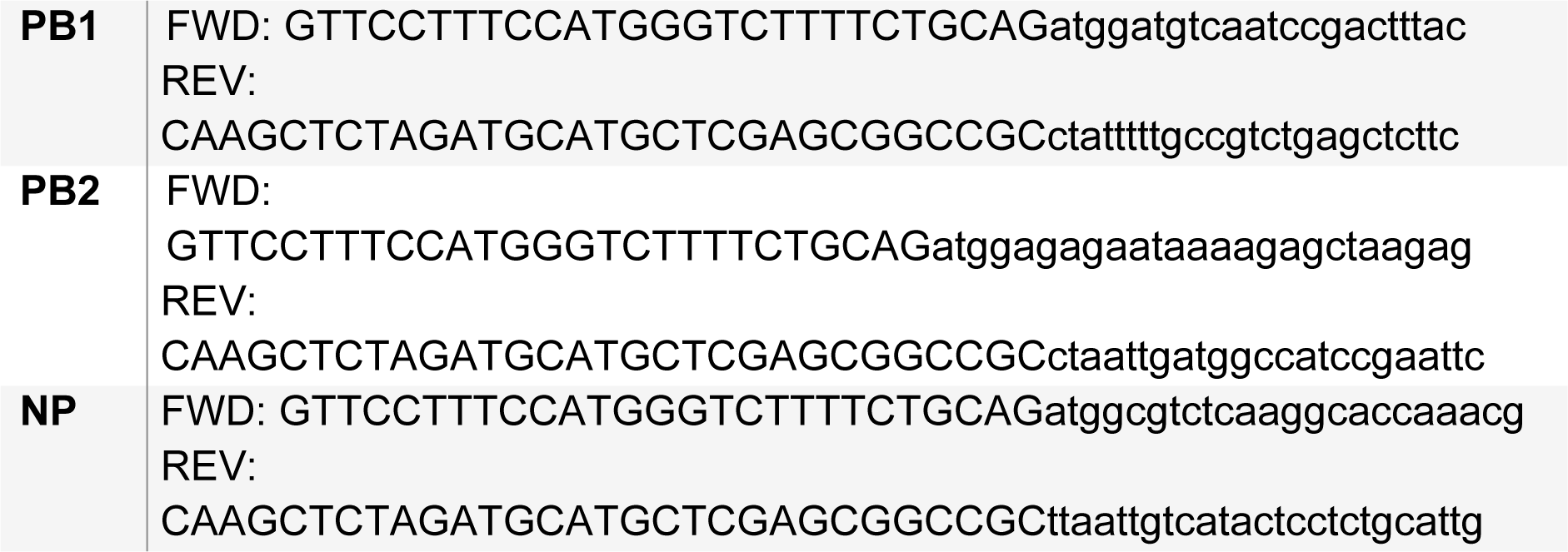
Primers used for cloning A/Caspian gull/Netherlands/1/2022Fragments. Overhangs for seamless cloning are indicated in upper-and gene-specific parts in lowercase letters

### Minigenome Reporter Assay

Human embryonic kidney cells (HEK293T) cells were seeded one day prior to transfection into a 6-well plate. 50-75% confluent cells were transfected with 0.5 µg of a plasmid encoding for a model viral RNA (vRNA) consisting of the firefly luciferase ORF flanked by the noncoding regions of segment 8 of influenza A virus under the control of a human RNA Polymerase I promoter and with 50 ng of a plasmid containing the ORF of the *Renilla* luciferase gene under the control of a CMV promoter as described previously^47^. To evaluate the differential activity of polymerases from different influenza A viruses, the cells were transfected with a set of plasmids encoding PB2, PB1, PA and NP genes the following amounts: 1 µg, 1 µg, 0.5 µg, 2 µg. Cells transfected with the plasmids expressing the model vRNA, *Renilla* luciferase and empty pCAGGS were also included as a negative control. The transfection mix consisted of 5 µg of each set of plasmids, 6.2 µL 2M CaCl_2_, 37.8 µL H_2_O and 50 µL 2x HBS (280mM NaCl, 10mM KCl, 1.5mM Na2HPO4, 12mM dextrose (glucose) and 50mM HEPES pH7.05), and was added to the cells after 5 minutes incubation at room temperature. 48 hours after transfection, luminescence was measured using the Dual-Luciferase Reporter Assay system (Promega) using a the Tecan Microplate Reader.

### Generating a consensus HA sequence of clade 2.3.4.4.b and an alignment

All available A/H5 HA nucleotide sequences and accompanying metadata were downloaded from the Global Initiative on Sharing All Influenza Data (GISAID) ^48^ and the Bacterial and Viral Bioinformatics Resource Center (BVBRC)^49^ databases on 13/06/2024. The sequences and metadata were pre-processed using the Pépinière jupyter notebook (https://github.com/epiv-lab/pepiniere). This included deduplication of sequences present in both datasets, identification and extraction of the start-stop open reading frame (ORF) corresponding to the longest ORF, and removal of sequences i) without metadata or ii) shorter than 90% of the average ORF length. Identical sequences were then grouped and only the earliest (by isolation date) representative was kept. ORFs were aligned using MAFFT v7.515^50^ and the alignment was trimmed to the start and stop codons of the majority of sequences. Trimmed sequences were again filtered to remove identical sequences, and the resulting 16507 sequences were realigned using MAFFT. A maximum-likelihood tree was generated using IQ-Tree2 v2.1.4^51^ with the GTR+F+R10 model (chosen by ModelFinder^52^) and 10000 UFboot bootstrap approximations^53^. The tree was midpoint rooted, annotated and visualized using iTOL^54^ in order to identify all sequences belonging to clade 2.3.4.4b. After translation, frame-shifted sequences were removed, and the remaining 6064 sequences were aligned using MAFFT. A plurality-based consensus as well as an interactive graph of amino acid frequencies observed at each position was generated using a custom Python (v3.9.17) script with the Biopython v1.79^55^, pandas v2.0.0^56^ and Plotly v5.15.0^57^ packages. The sequences were aligned with Clustal OMEGA^58^ and sequences similarities and secondary structure information were analysed with ESPRIPT3.0^59^.

### Statistical Analysis and Figures

Statistical differences between experimental groups were determined as described in the figure legends. P values of ≤0.05 were considered significant. Graphs and statistical tests were made with GraphPad Prism version 9. Protein structures PDB 4K66 and 4K67 were used for analysis and structures were analysed using PyMOL Version 2.5.8 and Chimera version 8.6.10^60^. The HA from the consensus sequence was modelled using Alphafold3^61^. Figures were prepared with Adobe Illustrator CC2019 and Adobe Photoshop CC2019.

## Results

### Recent clade 2.3.4.4b H5N1 virus attaches more abundantly to the human upper respiratory tract than clade 2.1.3.2 H5N1 virus

To check whether H5N1^2022^ virus showed amino acid changes in the receptor binding site (RBS) compared to other 2.3.4.4b viruses, we generated a consensus sequence of full-length HA of all available clade 2.3.4.4b HA sequences (sequences available at GISAID and BC-BRC) and aligned the HA sequences of H5N1^2022^ and H5N1^2005^ virus to it (Supplement Figure 1). We focused on the RBS since attachment to tissues can be impacted by amino acid substitution in this part of the HA^20^. The HA amino acid sequence of a HPAIV H5N1^2022^ differed in two amino acids positions compared to the 2.3.4.4b consensus sequence (D88G, M532I [H5 Numbering], Supplement Figure 1), both not located near the receptor binding site of the AlphaFold3 predicted consensus sequence HA structure (Supplement Figure 2A and Supplement Figure 2B). In comparison, H5N1^2005^ virus from clade 2.1.3.2 differed in several positions surrounding the receptor binding site (Supplement Figure 2C and Supplement Figure 2D).

Next, we determined the attachment pattern of H5N1^2022^, H5N1^2005^ and H3N2^2003^ viruses to the upper respiratory tract of humans, consisting of the nasal respiratory mucosa and olfactory mucosa, using virus histochemistry^62^. For this purpose, we generated recombinant viruses using the HA segment without the MBCS of the corresponding viruses^43^ alongside the other 7 segments of the mouse-adapted influenza A virus strain A/Puerto Rico/8/1934. In the upper respiratory tract of humans, attachment to the apical side of epithelial cells varied largely among the viruses (Table 3). The H5N1^2022^ virus attached to a moderate number of ciliated epithelial cells, while the H5N1^2005^ virus rarely attached to these cells, similar to what we observed previously^31^ (Table 3, Figure 1A). The seasonal H3N2^2003^ virus attached more abundantly to ciliated epithelial cells compared to both H5 viruses. Both H5N1^2022^ and H3N2^2003^ viruses attached to epithelial cells in submucosal glands, which was not detected for H5N1^2005^ virus. All three viruses attached to the olfactory mucosa and the Bowmans glands, although with differences in abundance as H3N2^2003^ virus attached most abundantly. The pattern of virus attachment to the human nasal respiratory mucosa (Figure 1A) showed a similar trend to that observed in primary human nasal epithelial cells (MucilAIR) (Figure 1B): abundant for H3N2^2003^, negative for H5N1^2005^, and moderate for H5N1^2022^ (Table 4).

**Figure 1.**
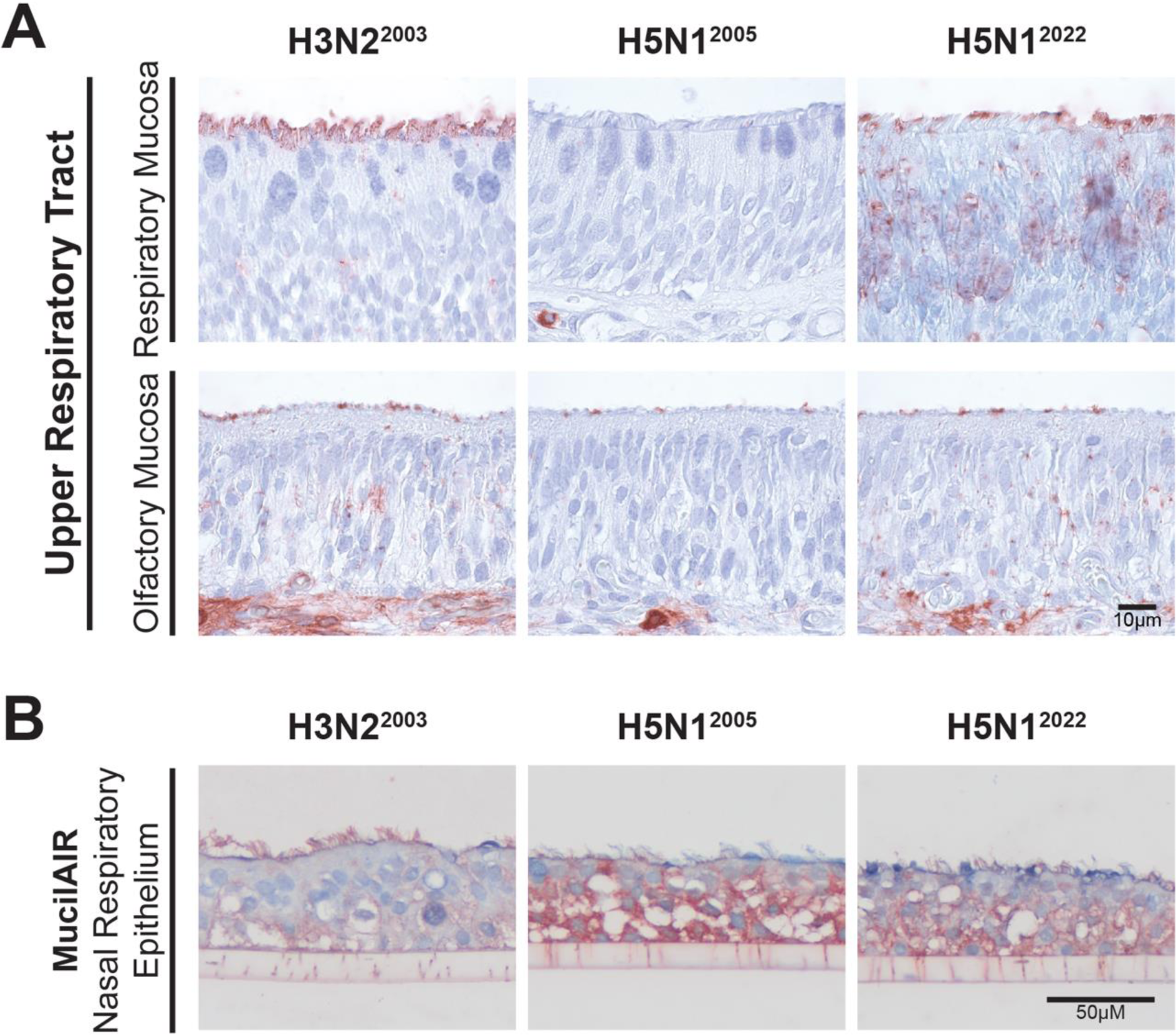
Virus attachment to the apical side of the human upper respiratory tract. Attachment of seasonal H3N2^2003^ virus and highly pathogenic avian influenza A viruses (H5N1^2005^ virus and H5N1^2022^ viruses) to (A) the respiratory and olfactory mucosa of the human nasal turbinate and to (B) human primary nasal epithelial cells (MucilAIR).

**Table 3.** Attachment of H3N2^2003^, H5N1^2005^ and H5N1^2022^ viruses to the apical side of the epithelium in human upper and lower respiratory tract tissues.

|  | Tissue | H3N2<sup>2003</sup> |  | H5N1<sup>2005</sup> |  | H5N1<sup>2022</sup> |  |
| --- | --- | --- | --- | --- | --- | --- | --- |
|  |  | Score | Predominant cell type | Score | Predominant cell type | Score | Predominant cell type |
| URT | Respiratory mucosa | ++ | cil+sg | - |  | + | cil+sg * |
|  | Olfactory mucosa | ++ | OM+bg | +/- | OM+bg | + | OM+bg* |
| LRT | Trachea | ++ | cil | +/- | cil+sg | + | cil+sg* |
|  | Bronchus | ++ | cil+sg | + | cil+sg | ++ | cil+sg*+gb |
|  | Bronchioli | ++ | cil | + | non-cil | ++ | cil+non-cil |
|  | Alveoli | ++ | Type I | +/- | Type II | + | Type (I), II, mph |

++, abundant (>50% cells positive). +, moderate (10%–50% cells positive); ±, scarce (<10% cells positive); –, negative (no attachment); OM, olfactory mucosa; Cil, ciliated epithelial cell; sg, submucosal gland; bg, bowmans gland; gb, goblet cell; type I type II refers to pneumocytes, mph, alveolar macrophages.

**Table 4.** Attachment of H3N2^2003^, H5N1^2005^ and H5N1^2022^ viruses to the apical side of the epithelium in human nasal respiratory epithelium and tracheal/bronchiolar epithelium.

| Tissue | H3N2<sup>2003</sup> |  | H5N1<sup>2005</sup> |  | H5N1<sup>2022</sup> |  |
| --- | --- | --- | --- | --- | --- | --- |
|  | Score | Predominant cell type | Score | Predominant cell type | Score | Predominant cell type |
| Nasal Respiratory Epithelium | ++ | cil | +/- | cil | + | cil |
| Tracheal/Bronchiolar Epithelium | ++ | cil | +/- | cil | + | cil |

++, abundant (>50% cells positive). +, moderate (10%–50% cells positive); ±, scarce (<10% cells positive); –, negative (no attachment); cil, ciliated epithelial cell.

In tissues of the human lower respiratory tract (trachea, bronchus, bronchiole and alveoli), H5N1^2022^ virus showed more abundant attachment to the apical side of ciliated and non-ciliated epithelial cells of the airways (trachea, bronchus and bronchiole) and type-I and type-II pneumocytes in the alveoli compared to H5N1^2005^ virus, which only attached to type-II pneumocytes and less abundantly to the other aforementioned cell types (Table 3, Figure 2A). However, H3N2^2003^ virus attached most abundantly to ciliated and non-ciliated epithelial cells in the airways and type-I pneumocytes, without abundant attachment to type-II pneumocytes in the alveoli(Table 3, Figure 2A). All viruses attached to tracheal/bronchiolar epithelial cells differentiated from airway organoids (AO) at air liquid interface (ALI), resembling the attachment pattern in human tracheal-bronchial epithelium (Table 4, Figure 2B). Taken together, the H5N1^2022^ virus attached more abundantly to both the upper and lower human respiratory tract compared to H5N1^2005^ virus, although not as extensively as seasonal H3N2^2003^ virus.

**Figure 2.**
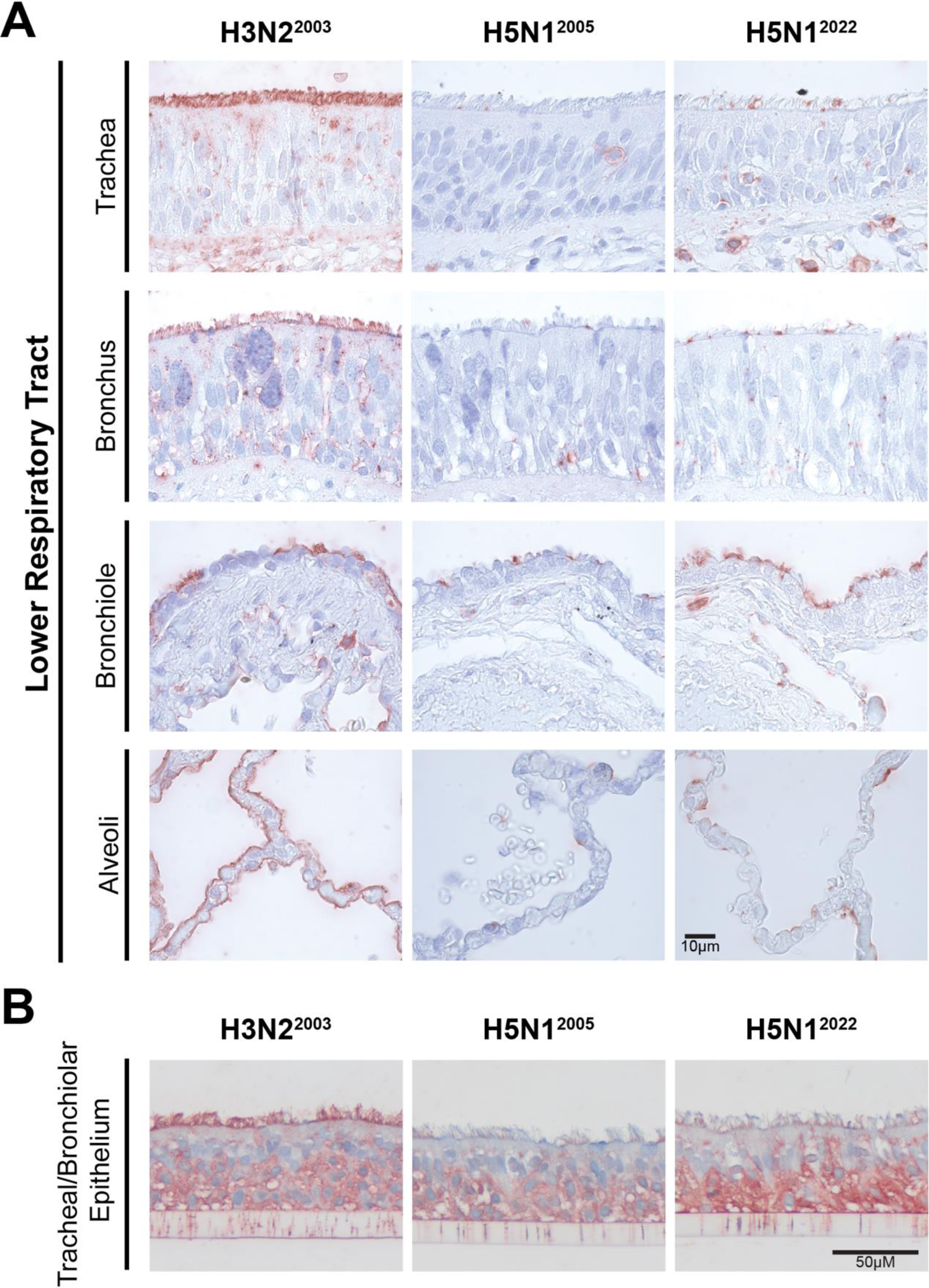
H5N1^2022^ attachment to the apical side of the human lower respiratory tract. Attachment of seasonal H3N2^2003^ virus and highly pathogenic avian influenza A viruses (H5N1^2005^ virus and H5N1^2022^ virus) to (A) trachea, bronchus, bronchiole and alveoli of the lower human respiratory tract and to (B) airway organoid-derived human tracheal/bronchiolar epithelium.

### H5N1 clade 2.3.4.4b virus replicates more efficiently in nasal respiratory and tracheal/bronchiolar epithelium compared to H5N1 virus from clade 2.1.3.2

Next, we investigated the replication efficiency of the HPAI H5N1^2022^ virus isolate in human primary nasal respiratory epithelium and in tracheal/bronchiolar epithelium differentiated from airway organoids (AO) at air liquid (ALI) interface. In the nasal respiratory epithelium cultures, both H5N1 viruses showed limited replication compared to the H3N2^2003^ virus (Figure 3A), although the H5N1^2022^ virus replicated to significantly higher titers compared to the H5N1^2005^ virus. In tracheal/bronchiolar epithelium cultures the H5N1^2022^ virus replicated to significantly higher titers than H5N1^2005^ virus and to similar titers as the H3N2^2003^ virus (Figure 3B).

**Figure 3.**
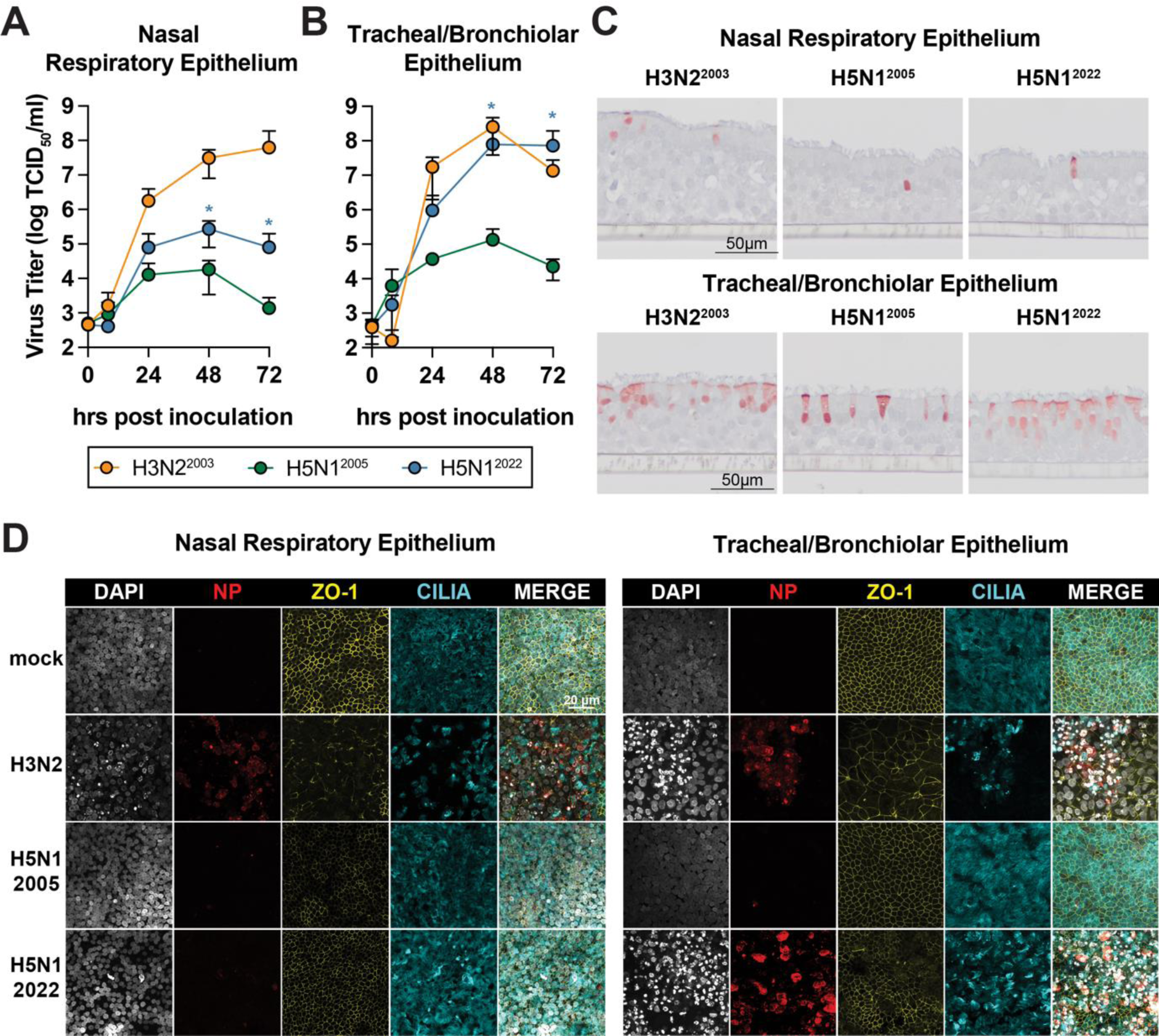
Replication of highly pathogenic avian influenza H5N1 viruses and H3N2 virus in human primary nasal respiratory epithelium and airway organoid derived tracheal/bronchiolar epithelium differentiated at air liquid interface. Replication kinetic of H3N2^2003^, H5N1^2005^ and H5N1^2022^ viruses in (A) nasal respiratory epithelium derived from MucilAIR (B) and in well-differentiated airway organoid-derived tracheal/bronchiolar epithelium at air–liquid interface (AO at ALI) with a multiplicity of infection 1. Differences in virus titers per time point between H5N1^2022^ and H5N1^2005^ were analyzed statistically with a student t-test. An asterisk indicates a statistically significant difference at p≤0.05. Replication kinetics were performed in technical duplicates in three independent experiments and data represented show the mean ± standard error of the mean SEM. (C) Detection of influenza A virus nucleoprotein (NP) by immunohistochemistry (IHC) in the nasal respiratory epithelium and tracheal/bronchiolar epithelium 24 h post-inoculation (hpi). (D) At 72 hpi, primary nasal respiratory epithelium cultures and tracheal-bronchiolar epithelium was stained for influenza A virus NP (red), the tight junction marker Zona Occludens-1 (ZO-1, yellow) and the cilia marker acetylated-α-tubulin (α-Tub, cyan). Nuclei were visualized with Hoechst. Maximum intensity projections of Z-stacks are displayed. Hrs, hours.

Immunohistochemical stainings of primary nasal respiratory epithelium and tracheal/bronchiolar epithelium with anti-nucleoprotein (NP) as primary antibody revealed that while H3N2^2003^ infected a plethora of cells, both H5 viruses infected only a limited number of ciliated epithelial cells in the nasal respiratory epithelium at 24 hours post inoculation (hpi) (Figure 3C). However, in the tracheal/bronchiolar epithelium, at 24 hpi the H5N1^2022^ and the H3N2^2003^ virus infected more cells compared to the H5N1^2005^ virus (Figure 3C). At 72 hpi, the epithelial cell layer infected with H5N1^2022^ and H3N2^2003^ virus showed evidence of epithelial necrosis and degeneration, based on its flattened appearance (epithelial attenuation), the presence of occasional giant epithelial cells (hypertrophy), and loss of cilia on the epithelial cells (Supplement Figure 3). The loss of cilia was verified through immunofluorescence staining for acetylated α-tubulin, showing that the H5N1^2022^ virus and H3N2^2003^ virus infected tracheal/bronchiolar cell were devoid of cilia at 72 hpi (Figure 3D). In addition, we observed loss of the tight junction marker Zona-Occludens-1 (ZO-1) in nasal respiratory epithelium and tracheal/bronchiolar epithelium in H3N2^2003^ virus infected cells. Despite abundant NP staining in H5N1^2022^ infected tracheal/bronchiolar epithelium, barely any loss of ZO-1 was observed. Collectively, these data show that the H5N1^2022^ virus replicates more efficiently in human nasal respiratory epithelium and tracheal/bronchiolar epithelial cells compared to the H5N1^2005^ virus. Especially in human tracheal/bronchiolar epithelial cells, H5N1^2022^ virus inoculation resulted in similar virus titers as H3N2^2003^ virus. Furthermore, there was evidence of loss of cilia, epithelial necrosis and degeneration in association with H5N1^2022^ virus replication in tracheal/bronchiolar epithelial cells.

### The polymerase activity of H5N1^2022^ virus is lower compared to seasonal H3N2^2003^ virus

To exclude that the increased replication of H5N1^2022^ compared to H5N1^2005^ in human respiratory epithelium is the result of an increased polymerase complex activity we compared the polymerase complex activity of H5N1^2022^, H5N1^2005^ and H3N2^2003^ viruses using a minigenome assay in HEK293T cells. The data suggest that the polymerase complex activity of H5N1^2022^ virus is not significantly different from that of H5N1^2005^ virus but significantly lower compared to H3N2^2003^ virus. That indicates that the polymerase activity of H5N1^2022^ virus likely does not contribute to the increased replication efficiency of H5N1^2022^ virus compared to H5N1^2005^ in nasal respiratory epithelium and tracheal/bronchiolar epithelium.

### H5N1^2022^ virus infection of human nasal respiratory and tracheal/bronchiolar epithelium induces a robust innate immune response

To characterize the cytokine responses in the supernatants of human nasal respiratory epithelium and tracheal/bronchiolar epithelium cultures, we performed a multibead cytokine assay. The H3N2^2003^ virus, induced a robust type-I-interferon (IFN) and type-III-IFN response consisting of IFN-α2 and IFN-β, IFN-λ1 and IFN-λ2/3, (Figure 5A and Figure 5B), as described before^38^. H5N1^2005^ virus infection only induced the secretion of IFN-α1 in nasal respiratory epithelium. In contrast, H5N1^2022^ virus induced a type-I-IFN and type-III-IFN response in both respiratory cultures. In both respiratory cultures, H5N1^2022^ and H5N1^2005^ virus inoculation resulted in the production of interferon-γ induced protein 10 (IP-10). Unlike H3N2^2003^ virus, infections with H5N1^2022^ and H5N1^2005^ viruses did not induce the secretion of Interleukin-6 (IL-6) or Tumor Necrosis Factor-alpha (TNF-a) in the nasal respiratory epithelium. In tracheal/bronchiolar epithelium both H5 viruses and H3N2^2003^ virus secreted IL-6. Taken together, H5N1^2022^ virus infection triggers a cytokine profile than differs from H5N1^2005^ virus but resembles that of H3N2^2003^ virus.

**Figure 4.**
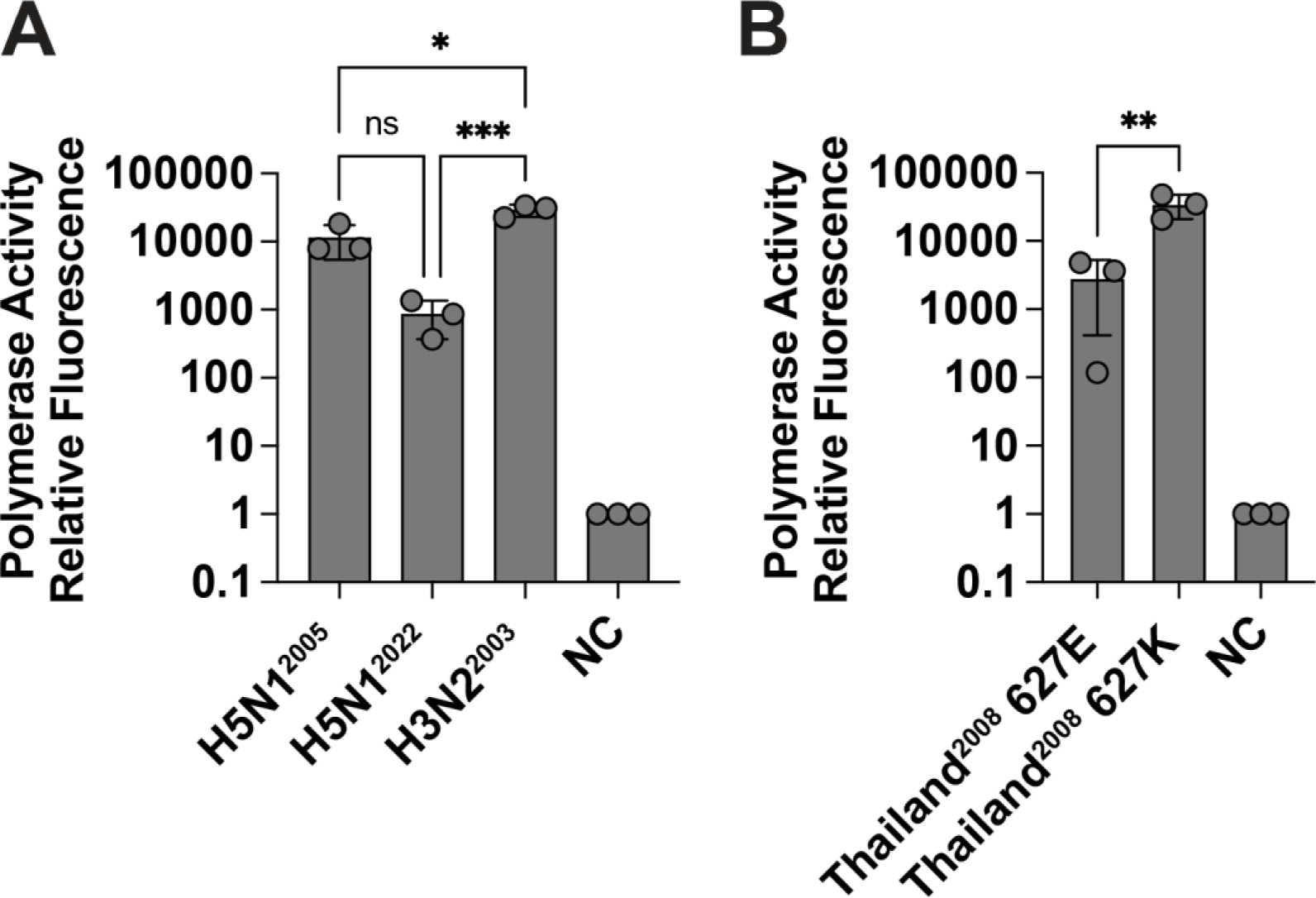
Polymerase complex activity of highly pathogenic avian influenza H5N1^2022^ virus from clade 2.3.4.4b. (A) Polymerase complex activity of H3N2^2003^, H5N1^2005^ and H5N1^2022^ viruses in PPI4 backbone 24 hours post transfection. (B) Polymerase complex activity of H5N1^2008^ virus harbouring either the PB2 amino acid 627E or 627K polymorphism at 24 hours post transfection pCAGGS backbone. For both graphs, values displayed are normalized to negative control (NC) which are cells transfected with the model vRNA, *Renilla* luciferase and empty plasmids. Data presented are averages of three independent experiments performed in technical triplicates ± standard deviation (SD). Statistical differences were determined using a One-Way-ANOVA with a Tukey’s multiple comparisons test. Asterisks indicate statistical significance: *, ns, non-significant; p < 0.05; **, p < 0.01; ***, p < 0.001; ****, P < 0.0001.

**Figure 5.**
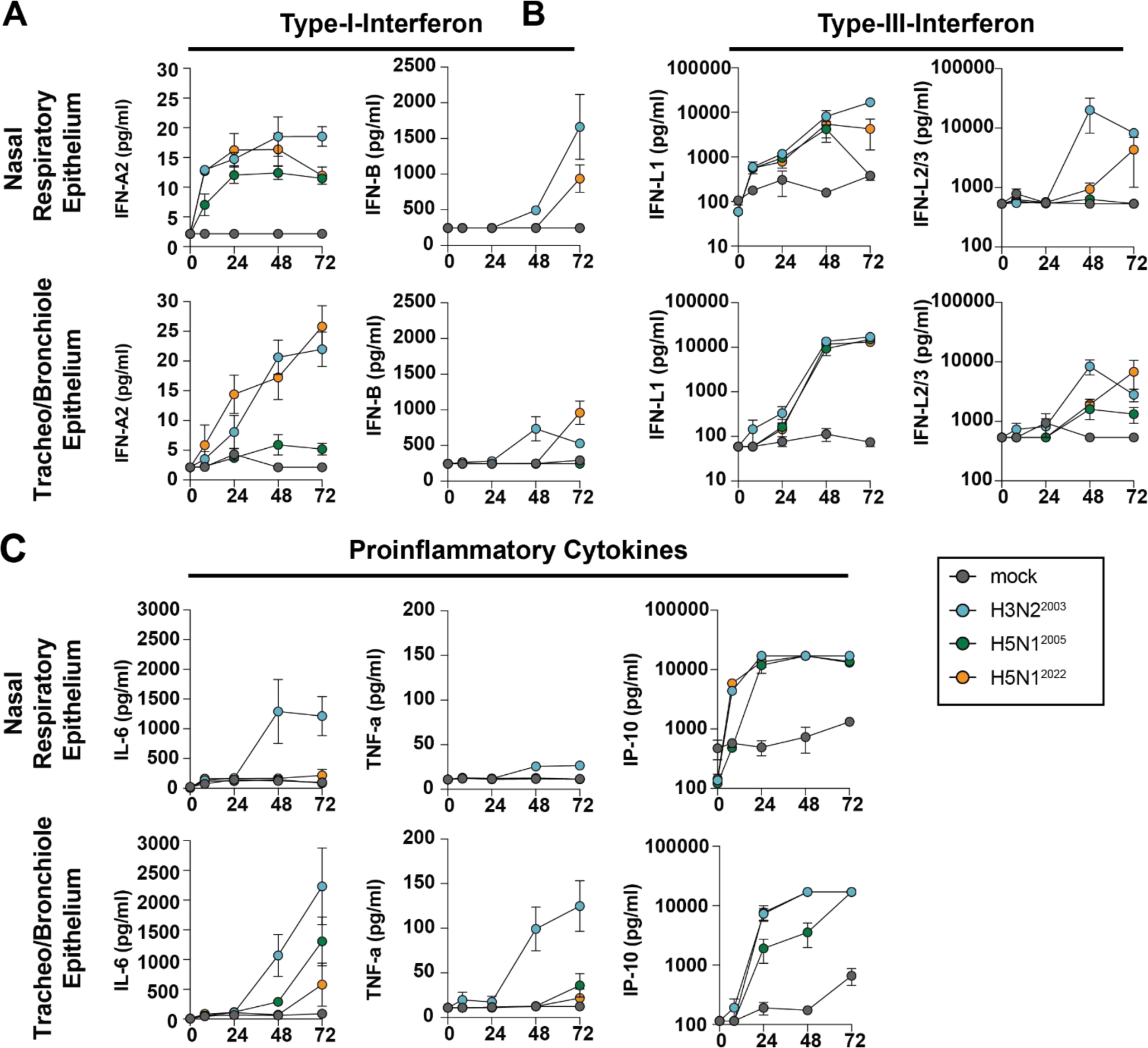
Innate immune responses in human nasal respiratory and tracheal/bronchiolar epithelium inoculated with H3N2^2003^, H5N1^2005^ or H5N1^2022^ viruses. Cultures of nasal respiratory epithelium and tracheal/bronchiolar epithelium were inoculated with H3N2^2003^, H5N1^2005^ or H5N1^2022^ virus at multiplicity of infection 1. Concentration of secreted (A) type-I-interferon (B) type-III-interferon and (C) proinflammatory cytokines in the supernatants of nasal respiratory epithelium and tracheal/bronchiolar epithelium were measured using a multibead cytokine assay. The displayed data are derived from three independent experiments performed in technical duplicates. The average ± standard error of mean (SEM) from every data point is depicted.

## Discussion

Here we show that a currently circulating clade 2.3.4.4.b H5N1 virus attached better to the human respiratory tract than a well-characterised 2.1.3.2 clade H5N1 virus. This difference likely contributed to more efficient replication and a more robust innate immune response in human respiratory epithelial cells.

Our data show abundant attachment of H5N1^2022^ virus to upper and lower respiratory tract tissues, which contrasts with H5N1^2005^ virus. This suggests that receptor binding repertoire of H5N1^2022^ virus has expanded to attach to receptors in the human upper and lower respiratory tracts. A recent observation revealed that a bovine H5N1 virus clade 2.3.4.4b isolate has the capacity to bind to α2–3 and α2–6 linked sialic acids^34^. However, several other studies using glycan arrays failed to show dual binding of either a bovine virus or avian virus clade 2.3.4.4b isolate to both sialic acid receptors ^63,64^. Although we observed more abundant binding of H5N1^2022^ virus, we cannot elucidate binding to specific glycans as VHC measures the attachment of virus to host cells directly. Our observed differences in the attachment pattern between the H5N1^2022^ and H5N1^2005^ viruses are likely the result of the observed amino acid substitutions close to the receptor-binding site, although the contribution of these individual substitutions remain to be established.

More abundant attachment of H5N1^2022^ virus compared to H5N1^2005^ virus in the human respiratory tract was associated with more efficient replication. The fact that the growth kinetics of H5N1^2022^ resembled seasonal H3N2^2003^ virus more closely than H5N1^2005^ virus could potentially explain recent studies in which inefficient airborne transmission is observed in ferrets, as well as the recent human cases^15,18,19,35,36,65^. The observed attachment pattern and/or the ability to replicate in human respiratory epithelial cells of H5N1^2022^ virus is not unique among avian influenza viruses. The airborne transmissible HPAI H5N1 virus infected more respiratory epithelium than H5N1^2005^, and showed airborne transmission between ferrets^22,26^. In addition, avian influenza viruses of the subtype H7N9 that emerged in China in 2013^66^ had similar abilities to attach to and replicate in human respiratory tract tissues. This was correlated with the ability of H7N9 viruses to be transmitted via air between ferrets and to infect humans^67–70^. Altogether, this suggests that the ability of influenza viruses to attach and replicate in respiratory epithelium of the airways in the upper and lower respiratory tract contributes to interspecies transmission.

Various factors such as polymerase complex activity and activation of innate immune responses contribute to the replication efficiency and dissemination of influenza A viruses *in vitro*^20^. The increased replication of H5N1^2022^ virus was not associated with an increased polymerase activity compared to H5N1^2005^ virus in human cells. However, we cannot exclude differences in the polymerase complex activity of H5N1^2022^ or H5N1^2005^ viruses in nasal respiratory epithelium or trachea/bronchiole epithelium is not known.

The replication of H5N1^2022^, H5N1^2005^ and H3N2^2002^ virus resulted in the induction of innate immune response. Overall, the type-I IFN, type-III IFN and I-6 responses of H5N1^2005^ virus were lower/delayed compared to H5N1^2022^ virus. Whether these differences are associated with the infection and replication efficiency, or intrinsic virus properties is not well understood.

Collectively, our data indicate that a H5N1 virus of clade 2.3.4.4b shows phenotypic changes in the attachment pattern, replication efficiency and immune response in the human respiratory tract compared to a well-studied H5N1 virus of clade 2.1.3.2. Although the exact factors required for efficient human infection and interspecies transmission are not fully understood, it is known that attachment and replication in the upper respiratory tract and tracheo-bronchial tree are important. The observed changes in attachment and replication efficiency warrants the urgency to prevent cross-species transmission to humans. This would include measures to prevent spill-over events to mammals and spread of H5N1 viruses in farmed mammals^71^. Improved surveillance systems covering different geographical locations as well as broader species coverage are necessary to be able to monitor changes in virus genome, allowing for monitoring virus adaptations to replication in mammals. Additionally, easy applicable methods such as VHC and replication kinetics in organoid-derived respiratory cultures allow for a relatively short turnaround time, which would enhance the real-time screening of new viruses or variants. These methods should be prioritized to assess the risk assessment of newly emerging H5N1 viruses.

## Acknowledgement

We thank Feline Benavides for technical assistance and for critically reading our manuscript. This study and D.V.R, M.I., T.K. and M.R are supported the European Union’s Horizon Europe FARM2FORK research and innovation programme Kappa Flu, grant number 101084171. L.B. is supported by The Netherlands Organization for Scientific Research (XS contract number OCENW.XS22.2.045) and by a grant 2023 from the European Society of Clinical Microbiology and Infectious Diseases (Europäische Gesellschaft für klinische Mikrobiologie und Infektionskrankheiten) (ESCMID). L.B is supported by fellowships from The Netherlands Organization for Scientific Research (VENI contract 09150162210154). M.P. is supported by the Coalition of Epidemic Preparedness Innovations (CEPI). R.D.D.V and L.L.A.V.D received funding from HERA EU4Health DURABLE. D.V.R. is supported by fellowships from The Netherlands Organization for Scientific Research (VIDI contract 91718308).

## Contributions

Conceptualization LB, LL, DvR

Investigation LB, LL, MI,

Formal Analysis LB, LL, MI, MS, MR

Resources LB, LL, MI, VC, MS, LvD, MP, WR, MF, RDdV, TK, DvR

Methodology LB, LL, MI, VC, MS, LvD, MP, WR, MF, TK

Supervision LB, TK, DVR

Visualization LB, LL, MI,

Writing Original Draft LB, LL, DvR

Writing-Reviewing all authors

Funding acquisition MR, RDdV, TK, DvR

