## Supplemental information for "A 2022 avian H5N1 influenza A virus from clade 2.3.4.4b attaches to and replicates better in human respiratory epithelium than a 2005 H5N1 virus from clade 2.3.2.1"

Bauer et al.

**Table of contents**

Supplementary Figure 1……………………………………………………………S2
Supplementary Figure 2……………………………………………………………S4
Supplementary Figure 3……………………………………………………………S6
References……………………………………………………………………………S7

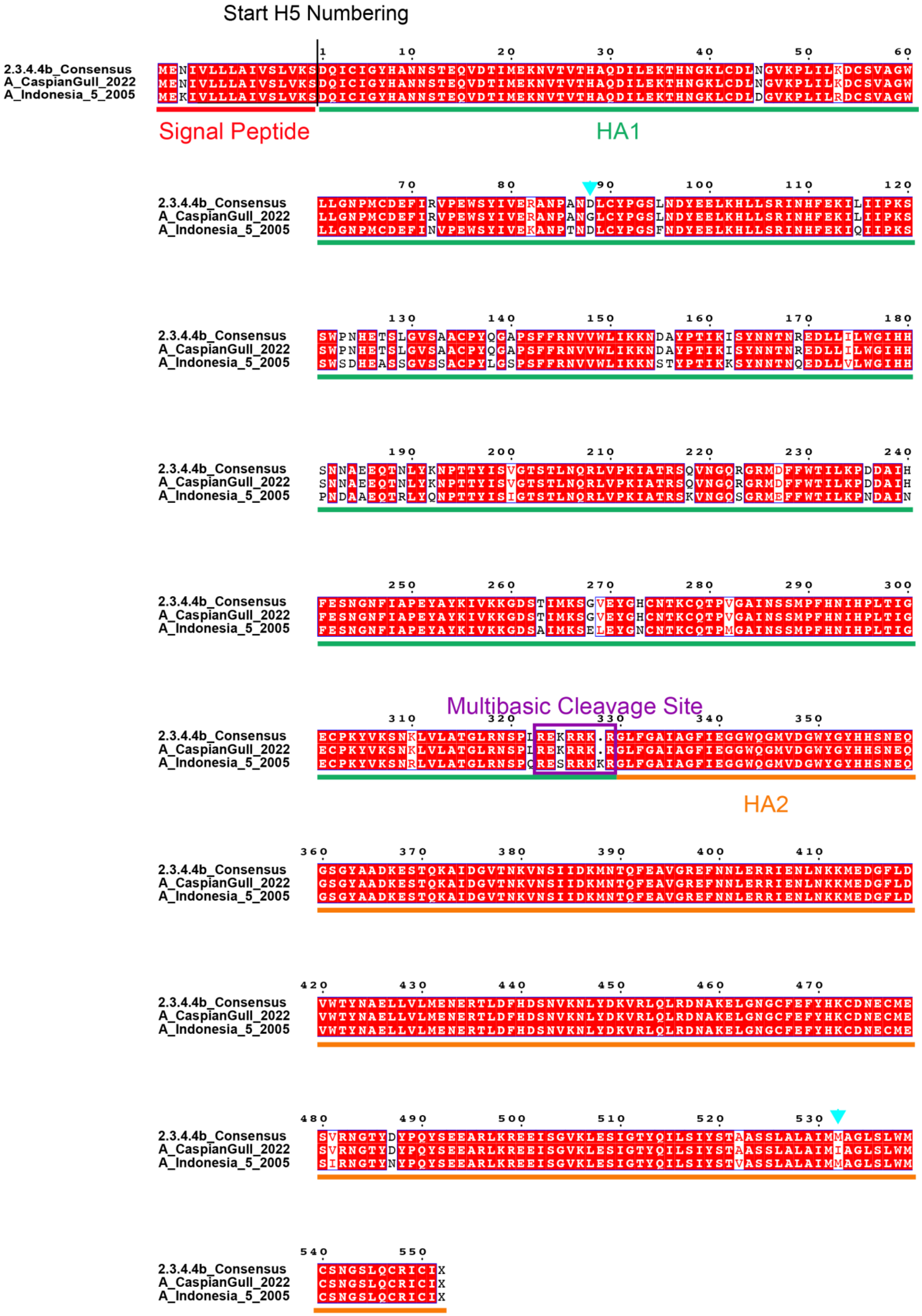

Start H5 Numbering
2.3.4.4b_Consensus
A_CaspianGull_2022
A_Indonesia_5_2005
Signal Peptide
HA1
Multibasic Cleavage Site
HA2

**Supplementary Figure 1. Sequence alignment of different H5N1 viruses**. A multiple sequence alignment of the consensus sequence of all available HA of clade 2.3.4.4b, A/Caspian gull/Netherlands/2022 ($H5N1^{2022}$) and A/Indonesia/5/2005 ($H5N1^{2005}$) was performed with Clustal OMEGA[1]. The ESPRIPT3.0[2] program was used to render sequence similarities. Differences in amino acids between 2.3.4.4b consensus H5 and $H5^{2022}$ amino acids are highlighted with cyan triangles.

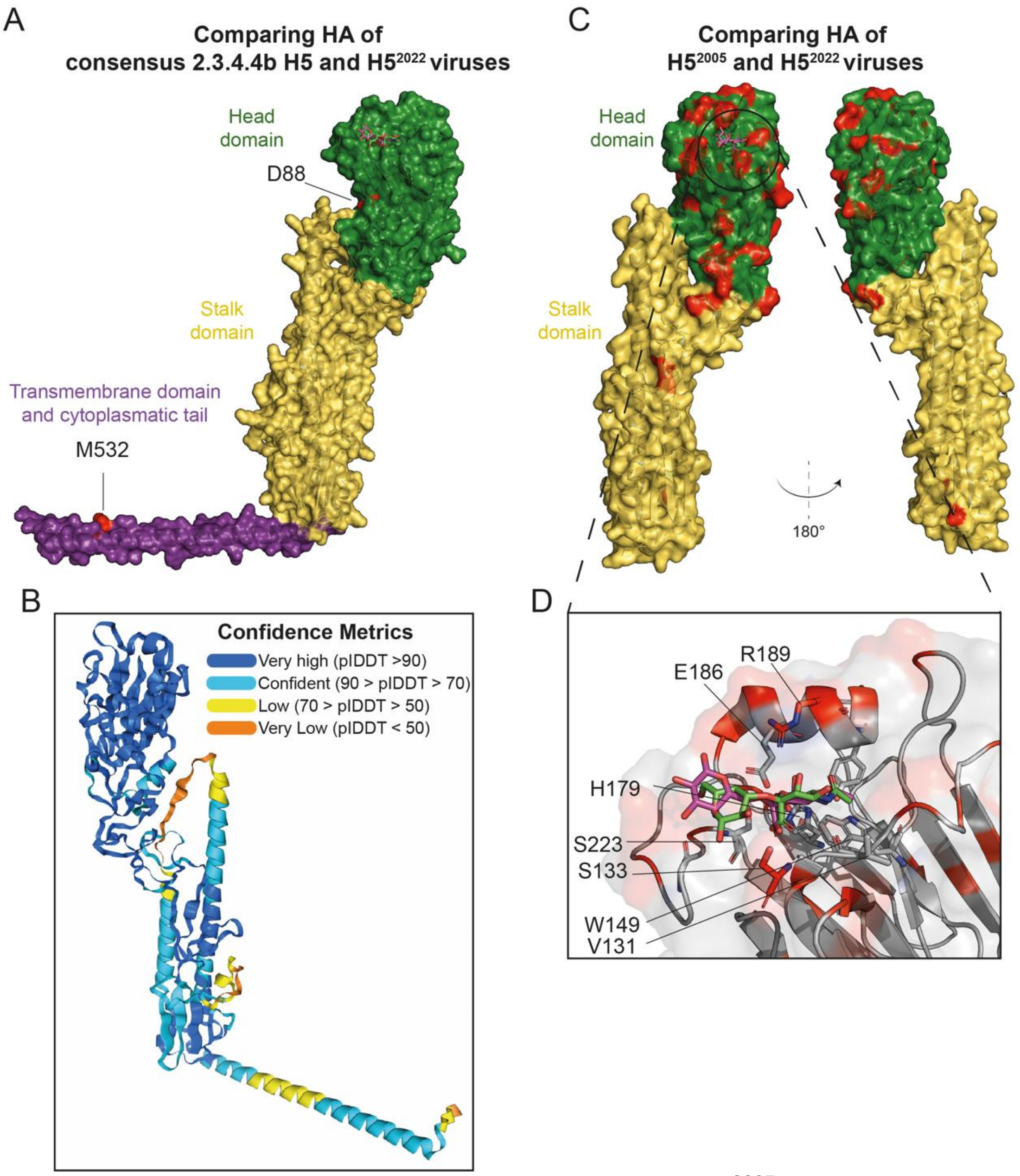


**Supplementary Figure 2. Structure of the H5N1$^{2005}$ virus hemagglutinin highlighting amino acid substitutions in the H5N1$^{2022}$ virus hemagglutinin.**
(A) Alphafold 3[3] prediction of consensus amino acids of clade 2.3.4.4b HA. Amino acid differences between the consensus HA of clade 2.3.4.4b compared to the HA of H5$^{2022}$ virus are highlighted in red. Modelled structure was overlayed with α2,3-linked sialic acid receptor in green (PDB: 4K66) and α2,6-linked SA receptor (PDB: 4K67) in purple. (B) Shows the corresponding confidence metric of the Alphafold 3 prediction[3] (C) The structure of H5N1$^{2005}$ virus hemagglutinin (HA) with the human α2,6-linked sialic acid receptor (PDB: 4K67) is displayed. Amino acid differences between the HA of H5N1$^{2022}$ and H5N1$^{2005}$ HA are highlighted in red. (B) Zoom in on the structure of the H5N1$^{2005}$ virus HA receptor binding site. Both the avian α2,3-linked sialic acid receptor (green; PDB: 4K66) and the human α2,6-linked sialic acid receptor (purple; PDB: 4K67) are displayed. The residues (H5 numbering) involved in the interaction with the receptors are shown as sticks. The residues that are mutated in the H5N1$^{2022}$ virus HA are highlighted in red.

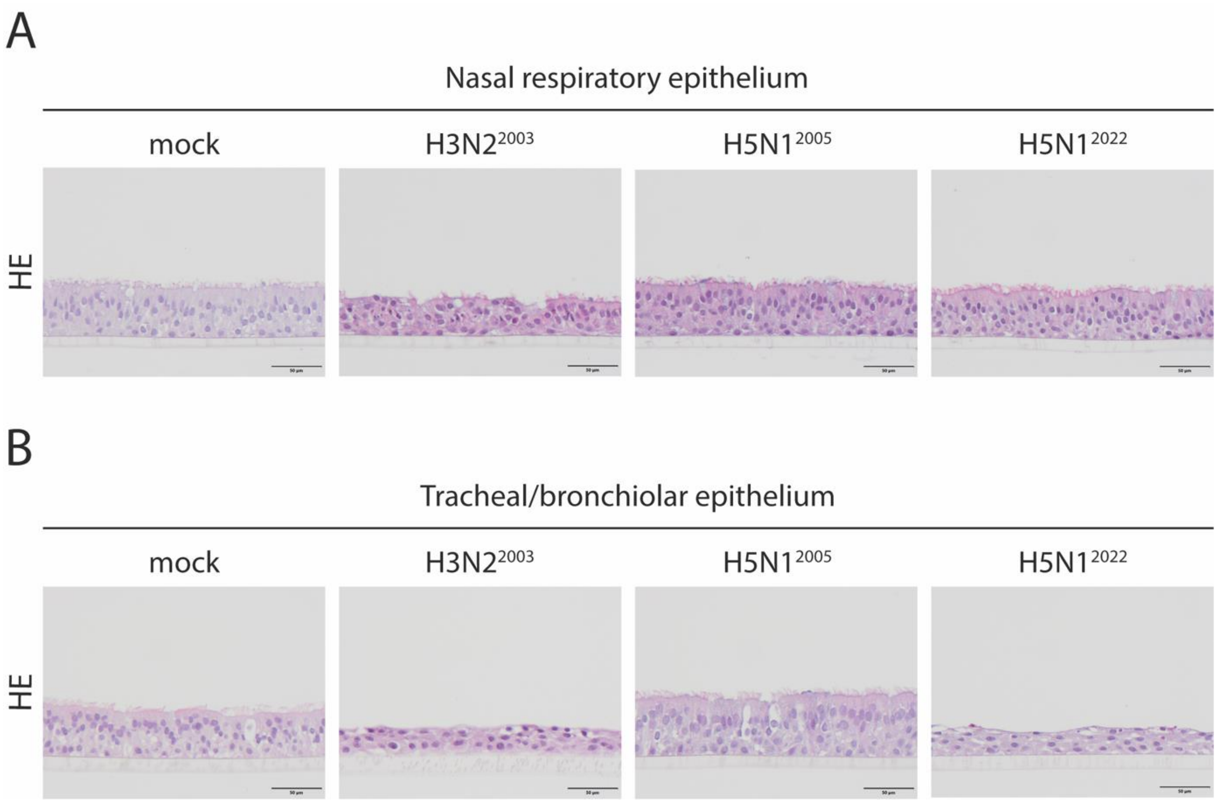


**Supplementary Figure 3. Hematoxylin and eosin (HE) staining of virus-infected MucilAir and ALI cultures**. HE staining of (A) human primary nasal epithelial cells (MucilAir) and (B) airway organoid-derived human trachea-bronchiole respiratory epithelial cells (ALI), mock and infected with H3N2, H5N1$^{2005}$ or H5N1$^{2022}$ with MOI 0.1 (MucilAir) or MOI 1 (ALI), at 72 hpi.
